# Abl signaling shapes the intrinsic fluctuations of actin to direct growth of a pioneer axon in *Drosophila*

**DOI:** 10.1101/511840

**Authors:** Akanni Clarke, Philip G. McQueen, Hsiao Yu Fang, Ramakrishnan Kannan, Victor Wang, Evan McCreedy, Stephen Wincovitch, Edward Giniger

## Abstract

The fundamental problem in axon growth and guidance is to understand how cytoplasmic signaling modulates the cytoskeleton to produce directed growth cone motility. We show here that the TSM1 pioneer axon of *Drosophila* extends by using Abl tyrosine kinase to shape the intrinsic fluctuations of a mass of accumulated actin in the distal axon. The actin mass fluctuates stochastically in length, but with a small, forward bias that drives the axon along its trajectory by promoting emergence of protrusions in leading intervals where actin accumulates, and collapse of protrusions in lagging intervals that actin has vacated. The actin mass is sculpted by Abl signaling, which probabilistically modulates its key parameters - its width and internal disorder - to drive its advance, while maintaining internal coherence. Comparison of TSM1 to other systems suggests that the mechanism we demonstrate here is apt to be common among pioneer axons in many organisms.

## Introduction

The process of axon guidance is central to patterning the nervous system during development. Understanding how axons pathfind to their correct targets requires that we know the mechanism by which guidance information in the environment controls the spatial organization and dynamics of the growth cone cytoskeleton to produce directed extension of the axon. It is widely accepted that in some cases broad, flat, growth cones extend the axon by harnessing the mechanochemical properties of large adhesive lamellipodia ^1,2^. There is reason to suspect, however, that other axons pathfind and extend in an entirely different way.

Cells in culture can switch mechanisms of cell motility, depending on the developmental context. For example, fibroblasts make large lamellipodia in 2-dimensional culture, yet assume spindle shaped morphologies and move faster in 3D environments ^3^. Mesenchymal cells use a slow moving adhesive style of growth in 2D culture, but switch to a fast ameboid style of growth in 3D conditions of low adhesion and strong confinement ^4^. Dendritic cells rely on integrins for 2D cell migration but 3D migration *in vivo* is protrusive and independent of integrin function ^5^. These examples demonstrate that the molecular components and mechanisms used by diverse cell types to move in 2D versus 3D environments are consistently and fundamentally different.

Much as cell motility can be accomplished in multiple ways, there is also evidence for a second mode of axon growth. It has been reported that filopodial protrusions alone can extend some axons. In these cases, it seems that rather than applying adhesive traction to lamellipodia to pull the growth cone forward, axon growth is instead accomplished by selective stabilization and dilation of individual filopodia ^6,7^. This filopodial style of growth has been proposed as a common mechanism for axon growth ^6,8^, but it is not clear how such a mechanism would work, or how axon guidance signaling molecules would regulate it.

To understand how signaling controls axon growth and guidance, there has been a sustained effort to find and characterize individual molecules with the goal of linking them to fundamental steps in the mechanism of axon extension and guidance. One such protein, the non-receptor Abelson Tyrosine Kinase (Abl) acts downstream of nearly all of the common, phylogenetically conserved families of axon guidance receptors (Bashaw et al., 2000; Crowner et al., 2003; Forsthoefel et al., 2005; Wills et al., 1999 Dajas-Bailador et al., 2008; Yu et al., 2001). Abl is critical for axon patterning (Grevengoed et al., 2001; Grevengoed et al., 2003; Liebel et al., 2003; Koleske et al., 1998; Moresco et al., 2005^13,14^, and acts in cooperation with a cohort of accessory factors that regulate the organization and dynamics of the actin cytoskeleton, including Enabled/VASP, Disabled, Abi, Trio, Rac and WAVE, among others ^15,16^. We have recently demonstrated that one key function of Abl signaling in axons is to balance linear extension of actin filaments with actin branching, through its ability to suppress the activity of a linear actin polymerizing factor (Enabled), and simultaneously stimulate an actin branching mechanism (WAVE/Scar and Arp 2,3, under control of Trio and Rac) ^17^. These roles place Abl at the key interface between extracellular signaling and intracellular modulation of the axonal cytoskeleton, though it remains unclear how these molecular interactions execute the cellular function of axon growth and guidance.

We therefore set out to use live imaging of a single pioneer neuron in its native environment to understand how Abl controls a pathfinding axon. We developed a method to image and computationally quantify both the neuronal morphology and the cytoskeletal dynamics of the TSM1 axon as it extends along its native trajectory in the intact *Drosophila* wing. We show here that Abl regulates the organization and redistribution of an accumulated actin bolus that locally controls filopodial morphogenesis to direct the extension and guidance of the TSM1 axon by locally regulating filopodial morphogenesis. Specifically, the distal portion of the TSM1 axon accumulates a bolus of actin that displays forward-biased spatial fluctuations, leading over time to its net advance during axon growth. Advance of the actin bolus, in turn, locally enhances the density of filopodial protrusions in the region where actin moves to, and the disassembly of protrusions in its wake, resulting over time in progressive advance of the filopodia-rich domain that defines the growth cone morphologically. The redistribution of actin is itself coordinated by the Abl tyrosine kinase signaling pathway, which modulates the spatial width of the actin distribution and also minimizes its disorder, allowing predictable translocation of the growth cone. Together, these data show how Abl probabilistically shapes the propagation of a leading actin mass that directs the growth and guidance of a pioneer axon extending in its native environment.

## Results

### Abl regulates TSM1 axon patterning

In light of the historical difficulty in observing cytoskeletal dynamics in growing axons in vivo ^19^, we have developed a system amenable to such inquiry by employing a pioneer sensory neuron of the *Drosophila* wing, called TSM1. During metamorphosis, the TSM1 axon extends in the space between the dorsal and ventral epithelia of the wing, pioneering the growth of the L1 nerve laterally from the wing margin before turning and extending proximally toward the wing hinge. The axon extends approximately 120*µ*m over a 9 - 12hr period along this trajectory before it fasciculates with the L3 nerve at the L1-L3 junction just distal to the GSR neuron (Figure 1A and 1B) ^20^. It has been shown previously that axons in cultured wing disc explants are robust and pathfind faithfully ^20,21^. Furthermore, the wing anatomy allows for an unobstructed view of TSM1 development. Therefore, TSM1 provides an excellent opportunity to observe the mechanism of axonal growth and guidance in detail as the growth cone pathfinds through its native environment.

**Figure 1:**
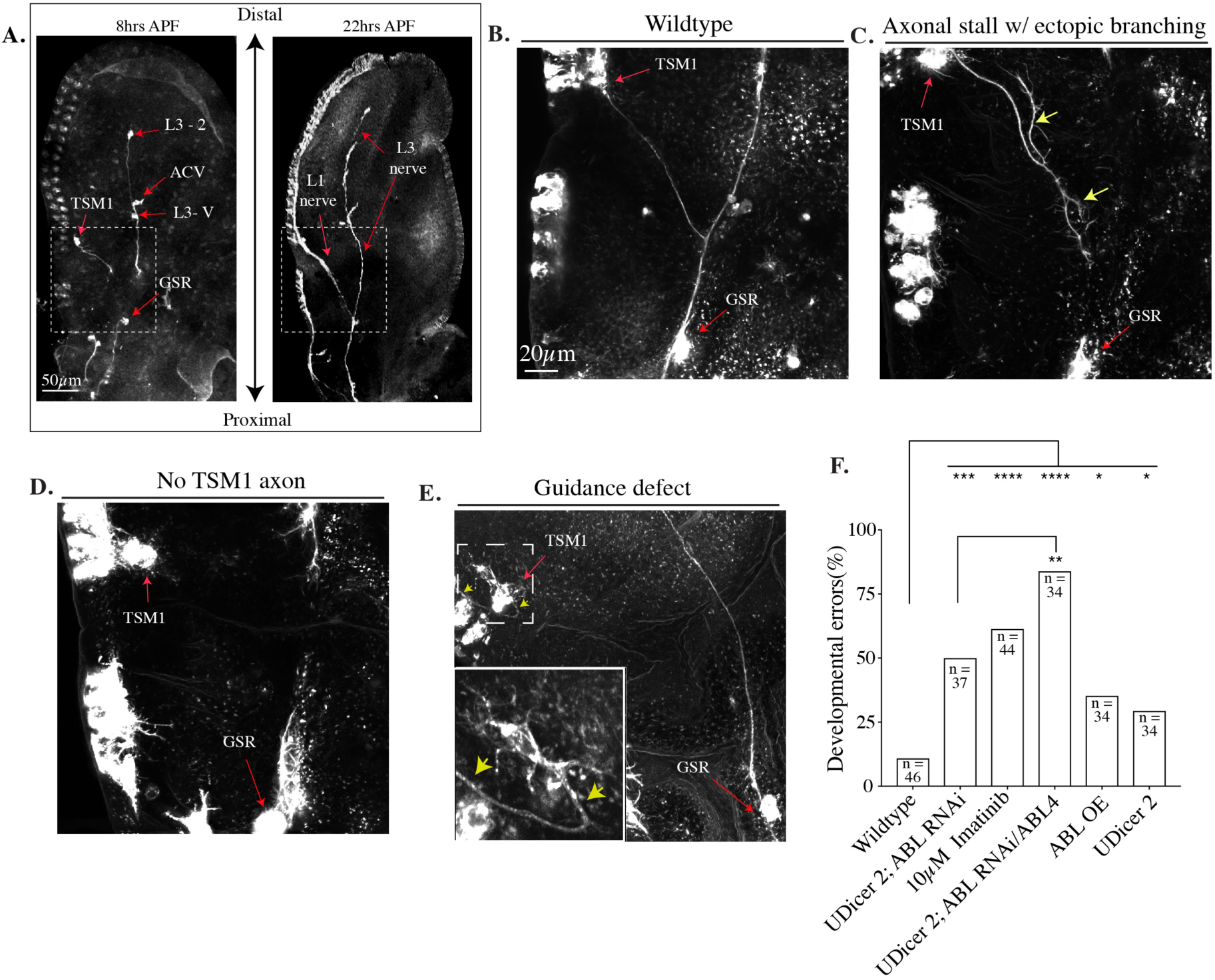
Abl regulates TSM1 axon guidance and extension. Early pupal wing imaginal discs at different developmental stages. A) 10x Micrographs of wing discs immunostained for the neuronal membrane (anti-HRP) and microtubules (mAb22c10). TSM1 and the L1 nerve, as well as GSR and other neurons comprising the developing L3 nerve are highlighted. B-E) 25x micrographs of terminal TSM1 axon phenotypes at ∼24h APF. B) Wildtype TSM1 developmental pattern, (C–E) Examples of observed developmental errors. Yellow arrows in C and E highlight aberrant TSM1 axonal phenotypes. The specific phenotypes depicted were not restricted to individual genotypes (C) neur-GAL4; UDicer2; Abl RNAi/Abl4, (D, E) neur-GAL4; UDicer2; Abl RNAi. F) Histogram plotting the frequency of terminal TSM1 developmental errors in the indicated genotypes. * = P < 0.05, ** = P < 0.01, *** = P < 0.005, **** = P < 0.0001; Fisher’s exact T-test.

We first verified that the Abl tyrosine kinase is required for growth and guidance of the TSM1 axon, just as it is required for many other axons in the CNS and PNS of *Drosophila*, and of vertebrates ^9–12^. We found that perturbing Abl activity or expression caused aberrant TSM1 axon growth and guidance phenotypes including instances where axonal growth stalled (Figure 1C), no axon extended from the cell body (Figure 1D), the axon misrouted along its trajectory (Figure 1E), and/or the axon maintained aberrant collateral branches (Figure 1C). Specifically, reducing Abl expression by RNAi in the background of Dicer 2 overexpression (Abl KD) caused an increase in aberrant TSM1 axon growth and guidance phenotypes (UAS-Abl RNAi, 51% vs. control 11%; p < 0.0001, Figure 1F). The expressivity of TSM1 axon defects observed in Abl KD was enhanced by heterozygosity for an Abl mutant allele (83% with UAS-Abl RNAi/Abl4 vs. UAS-Abl RNAi, 50%; p = 0.002, Figure 1F), confirming the fidelity of the RNAi phenotype. Expression of Dicer2 alone also increased the frequency of subtle defects in TSM1 morphology (32% vs 11% in control, Figure 1F) but the spectrum of defects is significantly different from that in the presence of Abl RNAi (p = 0.0035, chi-square, Supplemental Figure 2) and overexpression of Dicer 2 alone did not affect cytoskeletal parameters measured in growing axons (see below). To discriminate the catalytic function of Abl kinase from its scaffolding role, we treated wings with Imatinib, a specific inhibitor of Abl Kinase activity, and found significant TSM1 axonal defects when compared to control (61% in 10*µ*M Imatinib vs. control 11%; p < 0.0001, Figure 1F). This result strongly suggests that kinase activity is critical, though it does not exclude an important function for scaffolding. We also observed axonal defects when Abl was overexpressed (Abl OE 35% vs. control 11%; p = 0.0123, Figure 1F). Taken together, these results demonstrate that precise tuning of Abl function is required for TSM1 axon extension and guidance.

**Figure 2:**
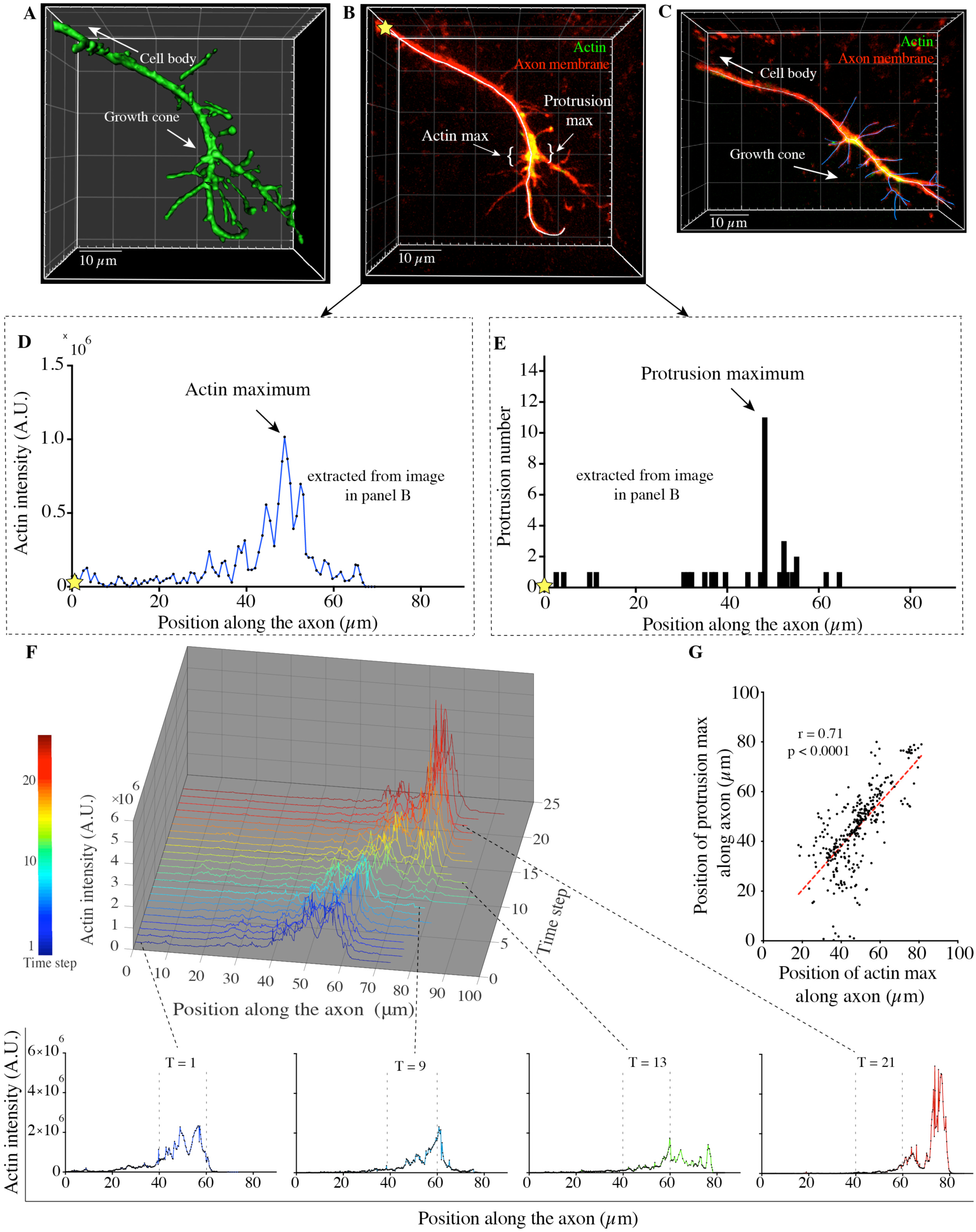
The TSM1 growth cone accumulates actin and is dominated by filopodial protrusions. The TSM1 axon was imaged in the intact, explanted wing. Image stacks were segmented and actin intensity was quantified along the axon shaft. (A) 3-dimensional rendering of the cell membrane, and (B) z-projected image of the distal axon of TSM1 from one typical video frame from wild type. In (B), membrane is in red, actin is in green, and the axon shaft has been traced in white. (See supplemental Figure 3 for unmerged color channels and pseudocolored actin channel). C) A fully segmented image of the distal TSM1 axon: protrusions are traced in blue and the axon shaft is traced in white. D) A representative actin distribution extracted from the lifeactGFP signal in the axon in 2B. The yellow star demarcates the origin of the actin profile. E) A representative distribution of protrusions extracted from the segmented axon in 2B. The yellow star demarcates the origin of the actin profile. F) A 3-dimensional plot of sequential actin distributions extracted over time from a single TSM1 trajectory. Profiles were extracted from images captured every 3 minutes. Individual distributions are highlighted to show actin “inchworming”. G) Scatter plot of the positions of the protrusion maximum vs the actin maximum along the axon. n =338 timepoints from 14 individual trajectories; Pearson r correlation coefficient; paired t test.

### Actin accumulates in the filopodial TSM1 growth cone

We next live - imaged TSM1 axon extension in explanted early-pupal wing imaginal discs by spinning disc confocal microscopy, with simultaneous visualization of the axonal membrane, using neuron-specific expression of CD4 Tandem-Tomato, and intracellular actin, using LifeactGFP (which labels both F-and G-actin ^22^) in wild type, Abl KD and Abl OE backgrounds, respectively. Multiplexed Z-stacks were collected every 3 minutes for 1.5 hours from fourteen trajectories of each genotype. Axon morphology was traced stereoscopically in 3-dimensions, and the intensity of the LifeactGFP signal was quantified along the axon shaft, from the base of the axon to its distal tip. We focus on actin organization both because of its critical role in essentially all forms of motility, and more specifically because the most extensively characterized output of Abl signaling is its modulation of actin regulators. Control experiments showed that imaging of developing transgenic wings in culture does not disturb the trajectory of the axon, or the ability of these axons to reach the L1-L3 junction and fasciculate with the L3 nerve (data not shown).

Live-imaging revealed consistent, concerted advance of a zone of enhanced filopodial density in the distal axon, together with an intra-axonal mass of accumulated actin, as the axon extended. In all imaged timepoints, filopodia and transient axonal branches (referred to collectively as protrusions) were the dominant morphological features of the axon. Large lamellipodia were seen only very rarely (<1% of time points) (Figure 2A - C). While the length and density of protrusions were highly variable along the axon, a region with enhanced density of protrusions was almost always observed in the distal portion of the axon (Figure 2B and E). Similarly, we invariably observed a local accumulation of high actin intensity within the distal axon (Figure 2B – D and Supplemental Figure 3), and this accumulation of actin advanced as the axon grew (Figure 2G and 3E and Supplemental movie 1). We refer to the curve that reports the relative actin intensity as a function of its position along the axon as the “actin distribution” throughout this manuscript (see, for example Fig 2D). Note that standard confocal imaging methods can only record the dynamics of this bulk actin distribution, which incorporates actin transport, diffusion, polymerization/de-polymerization, branching/de-branching, etc., it does not resolve the motions of individual actin molecules. Actin accumulation in the distal axon was also observed using other actin markers, including F-tractin Tandem-tomato, which selectively labels F-actin, and actin-GFP, whereas a volume marker (cytoplasmic eGFP) showed little or no distal accumulation (Supplemental Figure 4). Time-lapse image sequences suggested that the mass of accumulated actin and the zone of enhanced protrusion density seemed to advance in concert as the axon grew. To assess quantitatively the relationship between accumulated actin and protrusion density along the axon, we first performed a sliding window analysis to determine the position of the maximum value for each of these properties along the TSM1 axon in each image. Quantification confirmed that the interval with the highest actin level (actin maximum) and the interval with the highest protrusion density (protrusion maximum) were strongly correlated in position along the axon, (Pearson r = 0.71; p < 0.0001) (Figure 2F), and this held true across a range of window sizes (range 1 – 10*µ*m). Furthermore, the correlation of actin intensity values with local protrusion count was not restricted to the distal co-concentration of actin and protrusions. Global comparison affirmed the relationship between actin intensity and local protrusion count along the entire axon (Pearson r correlation = 0.60; p < 0.0001). This relationship was also consistent when we compared the lengths of protrusions, rather than the count, with the distribution of actin intensity (Pearson r correlation = 0.45; p < 0.0001). These data demonstrate that the length and number of axonal protrusions at particular positions along the axon shaft are strongly correlated to the local amount of accumulated actin.

**Figure 3:**
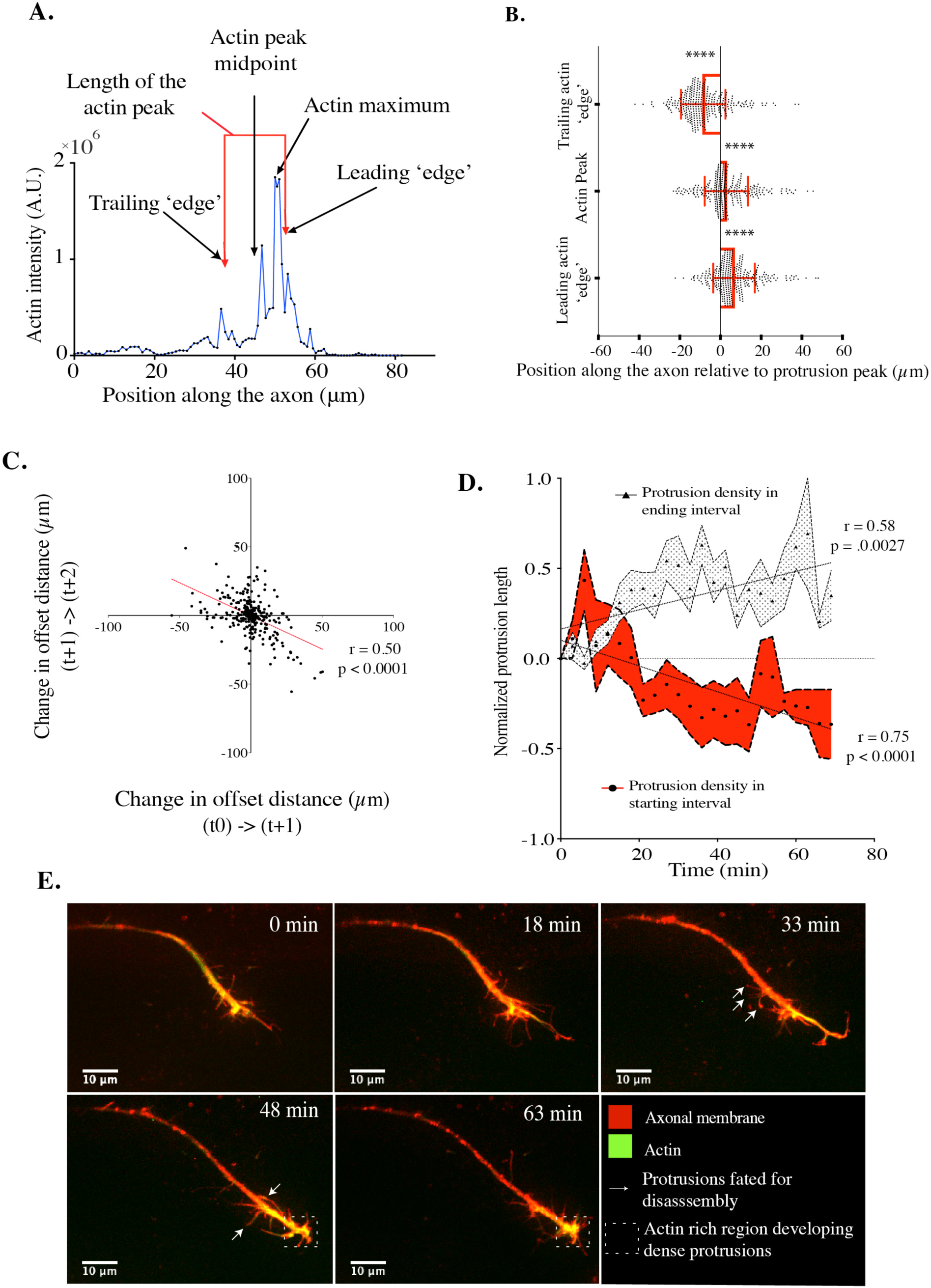
Actin leads and predicts the advance of the axonal protrusion density. A) Attributes of a representative actin distribution are annotated to demarcate the length, the maximum and the midpoint of the actin peak, as well as the nominal leading and trailing boundaries of the distribution (the leading and trailing square root of the 2^nd^ moment of the distribution, respectively; see text). B) The maximum of the actin distribution, as well as the leading and trailing ‘edges’ of the distributions are plotted relative to the position of the maximum protrusion density along the axon, which is set at the origin. Significance of the offset of these actin landmark positions relative to protrusion density in each respective timepoint was determined by two-tailed paired t-test. n = 338 timepoints from 14 individual trajectories, **** = p < 0.0001) Error bars are mean±SD. C) A scatter plot of the change in the offset distance between actin and protrusion maxima in a single time-step vs the change in the offset distance in the following time step. n = 338 timepoints from 14 individual trajectories; Pearson r. D) Average protrusion length over time is plotted for a 10μm interval centered on the midpoint of the actin distribution at the start of an actin translocation event (red swath), and for the interval centered 20μm distal to that start point (black and white swath). Protrusion length for each 20*µ*m actin midpoint translocation event analyzed was normalized to the summed protrusion length at t = 0. n = 128 time points from 7 individual trajectories; mean±SEM for each time point is plotted over time; Pearson r correlation coefficient. E) Gallery of images showing a time-course of TSM1 axon extension and actin advance (lifeactGFP, green; and CD4 tandem tomato, red; representative of n = 14 independent trajectories). White arrows highlight protrusions fated to disassemble over time as actin advances; white box highlights zone fated to develop dense projections once it is occupied by actin.. See supplemental Figure 5 for timecourse with unmerged color channels and pseudocolored actin channel).

**Figure 4:**
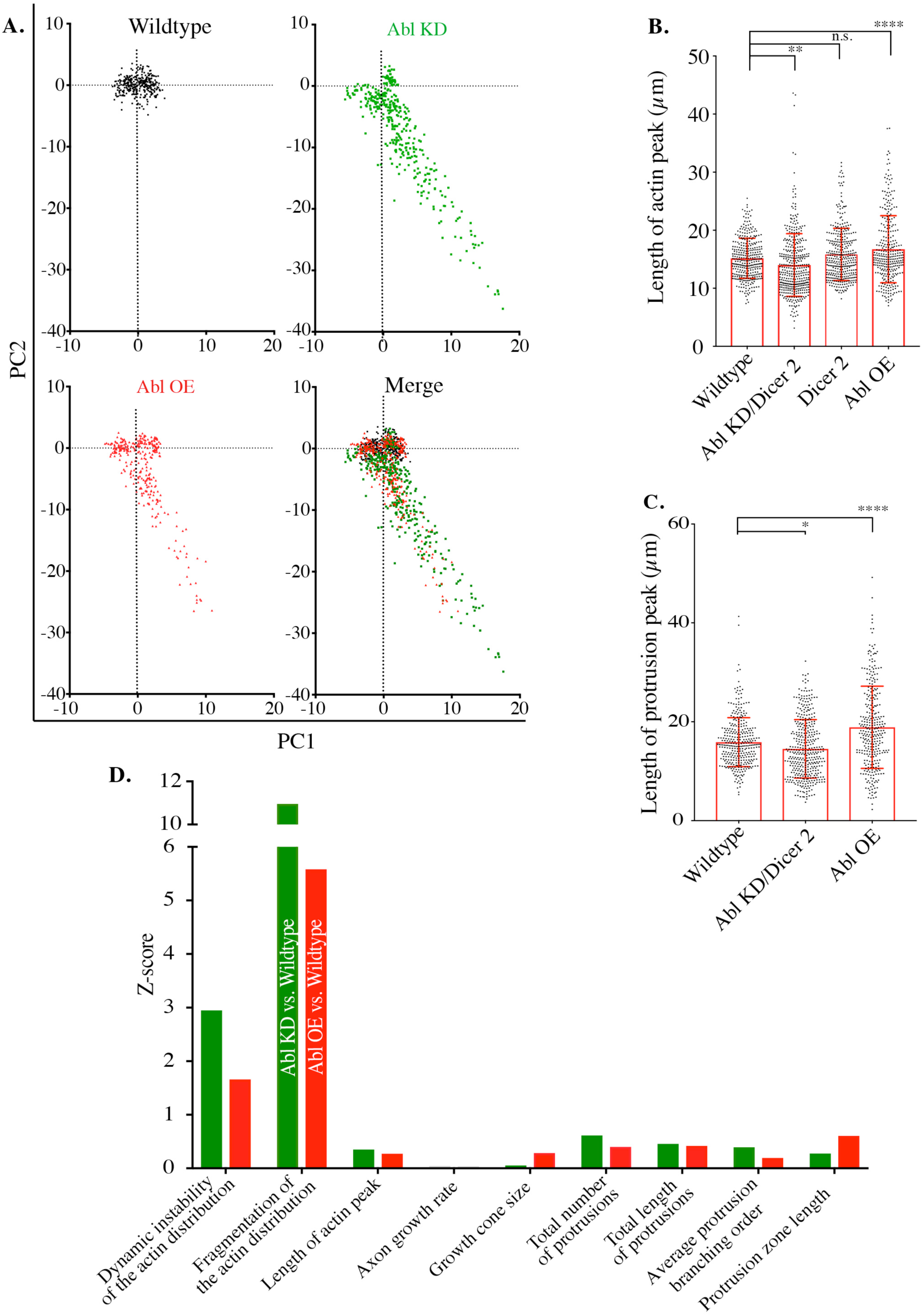
Abl perturbed TSM1 axons employ the same cell biology as wildtype. A) Data sets comprising 10 individual parameters extracted from Wildtype, Abl KD and Abl OE TSM1 axons were dimensionally reduced by PCA. Principal components 1 (PC 1) and 2 (PC 2) are plotted for each genotype and are representative of the parameter relationships within all pairwise combinations of the first 4 principal components, which account for > 70% of the total variance in the data sets (see text supplemental Figure 6). B) The length of the actin peak, as measured by the sum of the square root of the leading and trailing second moments about the actin peak in each time point (see methods), was quantified in wildtype, Abl KD, ABl OE and in the Dicer2 actin control. n > 300 time points from 14 trajectories for each genotype; significance determined by one-way ANOVA; **p < 0.01, ****p < 0.0001 n.s. = non-significant. C) The length of the protrusion peak, as measured by the square root of the second moment about the peak in protrusion density in each time point (see methods), was quantified in wildtype, Abl KD and ABl OE. n > 300 time points from 14 trajectories for each genotype; significance determined by one-way ANOVA; *p < 0.05, ****p < 0.0001. D) Z-scores for actin and morphological parameters were calculated for Abl KD (green) and Abl OE (red) relative to wildtype.

### Advance of the accumulated actin distribution predicts the local growth and disassembly of axonal protrusions

While we observed a consistent spatial correlation between localization of actin intensity and protrusion density, further analysis revealed that the maxima of the actin accumulation and the protrusion density are, in fact, slightly offset, with the actin maximum systematically leading the maximum of the protrusion density as the axon grows (Figure 3). We aligned all 338-wildtype time-points by the protrusion maximum along the axon and plotted the position of the actin maximum, as well as the positions of the proximal and distal boundaries of the accumulated mass of actin (referred to here as the “actin peak”). Empirically, we found it effective to define the ‘edges’ of the actin peak as the positions defined by the leading-and trailing-square root of the second moment of the actin distribution about the actin peak (a measure akin to the standard deviation of the distribution; see below and materials and methods) (Figure 3A). We found that the actin peak enveloped the position of the maximum protrusion density: the leading ‘edge’ of the actin peak leads the protrusion maximum by 6.7*µ*m ± 0.56 (P<0.0001), and the trailing ‘edge’ follows the protrusion maximum by 8.5*µ*m ± 0.59 (P<0.0001), while the maximum of actin peak led the maximum of the protrusion density by an offset distance of 2.8*µ*m ± 0.58, (mean±SEM, p<0.0001) (Figure 3B). As the TSM1 axon grows at an average rate of 0.23 *µ*m/min (as measured by a variety of metrics; see Supplemental Figure 5), this mean offset length implies that the position of the actin peak predicts where the bulk of protrusions will be found 10-15 minutes later. Furthermore, the offset distance between the actin maximum and that of the protrusion density appears to be maintained actively. We found a strong negative correlation between the rate of change of the offset distance between these two positions in pairs of successive time-points (r=-0.50, p<0.0001) (Figure 3C), which suggests that when the spacing between the actin and protrusion maxima increases or decreases in any given time step, the gap size tends to be restored during the next time point. Taken together, these data provide evidence that the actin accumulation peak leads the advance of the axonal zone bearing the highest density of protrusions, with the offset distance between these two axonal features being a regulated aspect of axon growth.

**Figure 5:**
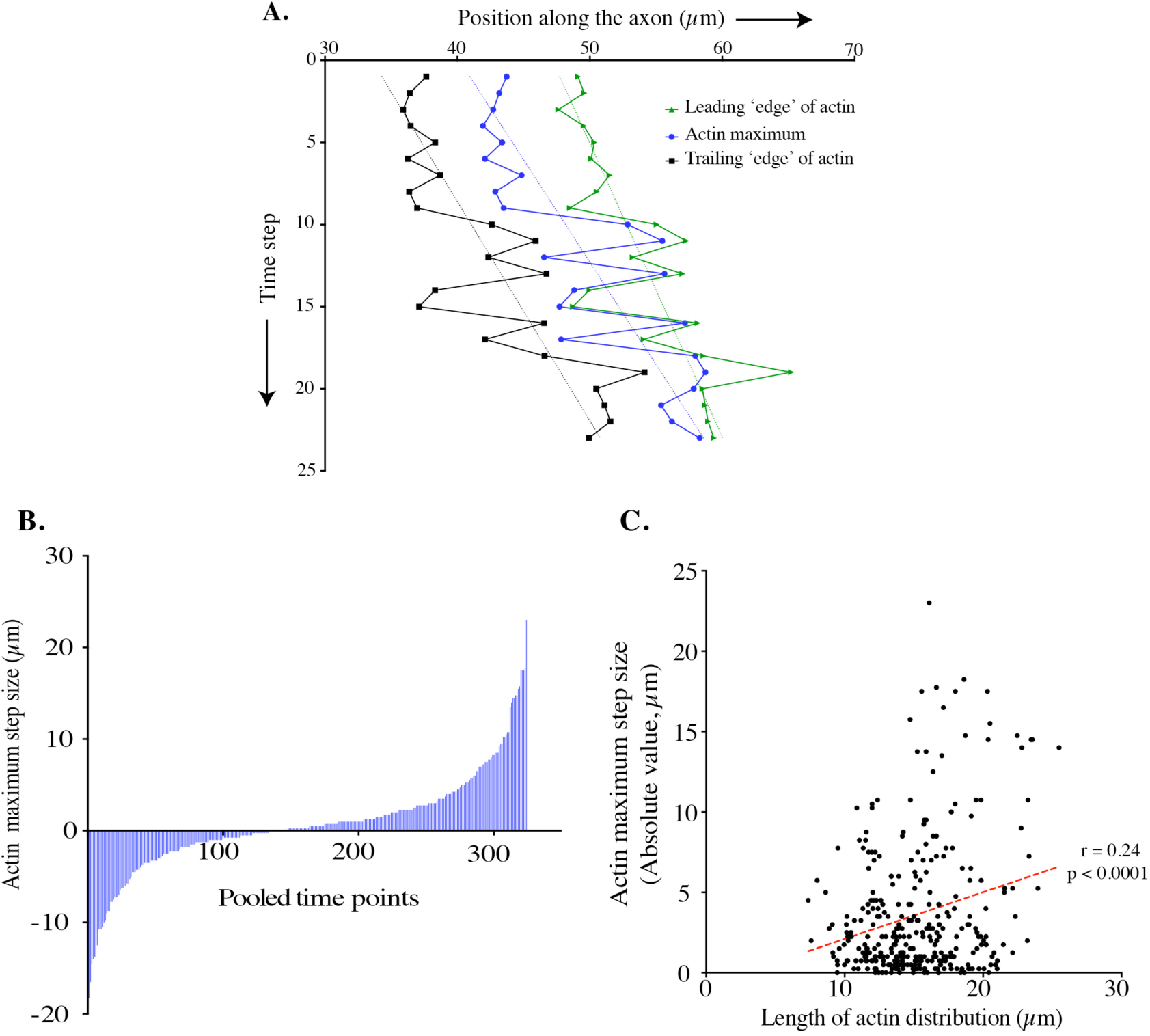
Biased stochastic fluctuations of accumulated actin move the distribution forward. A) A plot of the axonal positions of landmarks in the actin distribution from a single trajectory over-time. The actin maximum (blue), and the leading (green) and trailing (black) ‘edges’ of the actin distribution are highlighted. This plot is representative of n = 14 independent trajectories. B) A histogram of the length of excursions of the actin maximum in single time steps. n = 324 time steps from 14 individual wildtype trajectories. C) A scatter plot of the absolute value of the step size of the actin maximum versus the length of the actin distribution. n = 324 time steps from 14 individual wildtype trajectories; Pearson r correlation coefficient.

Based on these observations, we hypothesized that advance of the leading actin mass increases the density of protrusions at the more distal position the actin peak now occupies, and disassembly of protrusions at the position the actin has vacated. We therefore quantified the protrusion density in intervals associated with the starting and ending positions of the actin peak in 7 trajectories where the actin advanced 20*µ*m. Consistent with visual inspection of time lapse micrographs of TSM1 axon extension dynamics, we found that, after a lag phase of ∼15 min, summed protrusion length declined by 50% ± 0.1 (mean±SEM, p<0.001) in the region vacated by the actin peak, while at the new more distal position of the actin peak protrusion length was increased by 32% ± 0.1(mean±SEM, p<0.01), again with the actin peak on average reaching the final position prior to the protrusion maximum (Figure 3D, Figure 3E, Supplemental Figure 6, and Supplemental movie 1). Thus, the evolution of the actin distribution predicts where axonal protrusions will subsequently grow and retract along the axon.

**Figure 6:**
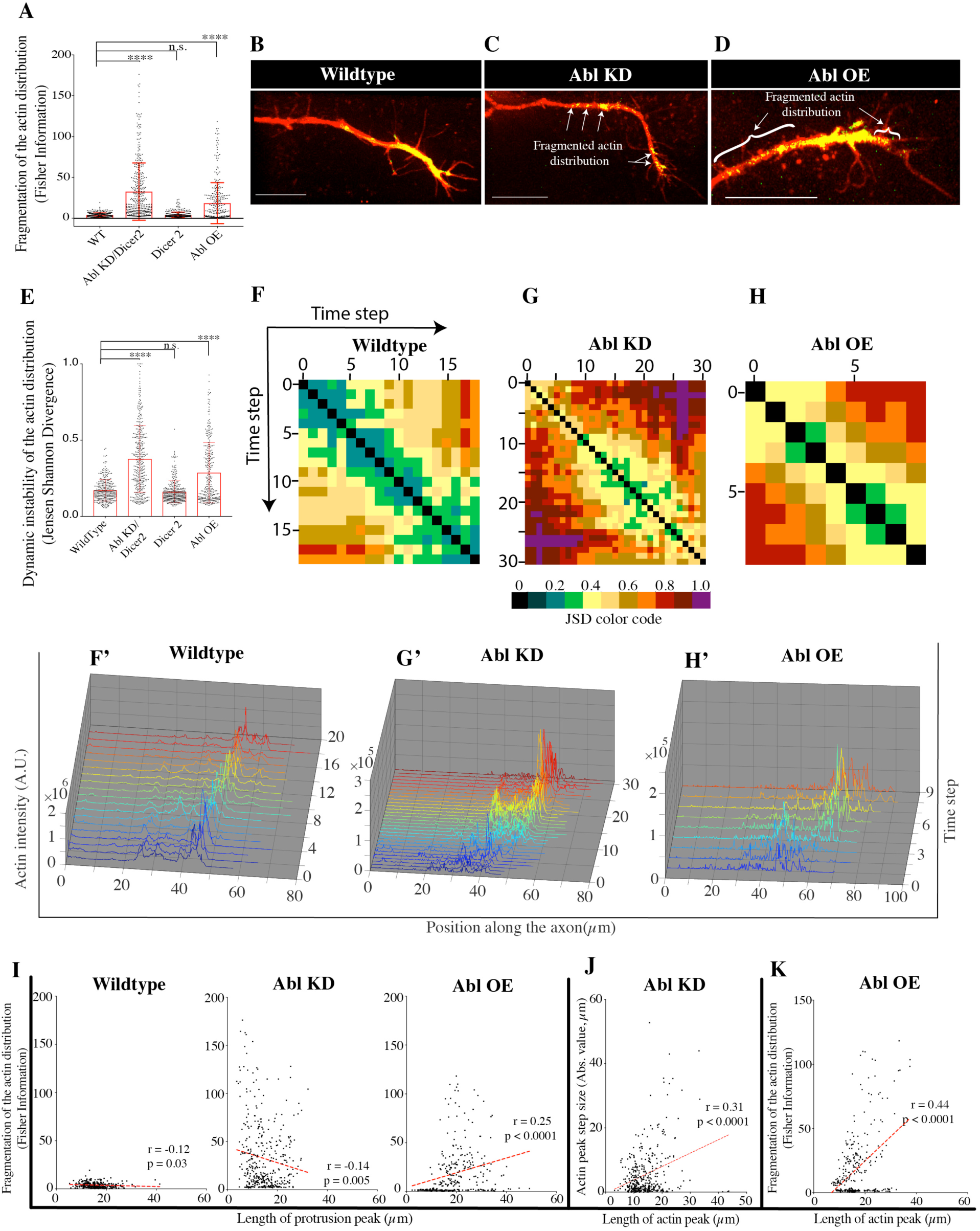
Abl perturbation fragments the actin distribution and disorganizes its advance. A) Fragmentation of the actin distribution, as calculated by the Fisher information content of each actin profile in each time point was quantified in the indicated genotypes. B - D) Representative examples of the mean actin fragmentation phenotype for each genotype. E) The dynamic instability of the actin distribution, as calculated by the Jensen-Shannon Divergence (JSD) of sequential pairs of actin profiles over time (see methods), was quantified in wildtype, Abl KD, ABl OE and in the Dicer2 actin control. For (A) and (E), n = 338 time points from 14 trajectories for each genotype; significance determined by one-way ANOVA; ****p < 0.0001; n.s. = non-significant; error bars indicate standard deviation. (F - H) The JSD of all pairwise combinations of the actin profiles from all time steps of one individual trajectory from each genotype were calculated, color coded, and are presented as matrices. In each case, the matrix shown represents a trajectory with a mean JSD value that is approximately average for that genotype. (F’ – H’) 3-D sequential actin distribution plots that correspond to the representative JSD matrices above. I) Scatter plots of the Fisher information versus the length of the protrusion peak for each respective genotype. J) Scatter plot of the absolute value of the actin peak step size versus the length of the actin peak. K) Scatter plot of the Fisher information versus the length of the actin peak. For I – K, n > 300 time point from 14 individual trajectories for each genotype; Pearson r correlation coefficient.

### TSM1 axons maintain parameter relationships in Abl KD and OE genetic backgrounds

The observations above suggest that the molecular mechanism that controls the structure and dynamics of the actin distribution is the key to regulating the morphological development and extension of the TSM1 axon. We therefore imaged TSM1 axon extension in Abl KD and Abl OE genetic backgrounds, computationally measured a broad set of parameters that capture axonal morphology and intra-axonal actin organization and compared these data sets to those from wildtype TSM1 trajectories. Among the parameters we investigated were morphological features, including the number, length, and branch order of protrusions, and the 3-dimensional volume of substratum explored by protrusions from the growth cone (here called growth cone volume), as well as measures of the actin distribution, including the length, organization, and speed of advance of the peak of actin accumulation (see Table 1 and Materials and Methods for a complete listing of imaged features and their definitions).

**Table 1:** Measured parameters of the TSM1 growth cone

| **Measured parameters of the TSM1 growth cone** |
| --- |
| **Total number of protrusions** |
| **Total length of protrusions** |
| **Average protrusion branching order** |
| **Growth cone volume** |
| **Length of growth cone protrusion peak**<br>(square root of the second moment about the protrusion peak) |
| **Length of the actin peak**<br>(square root of the second moment about the actin peak) |
| **Offset between position of actin maximum vs protrusion maximum** |
| **Axon extension rate:**<br>**Rate of change of position of the midpoint of actin peak** |
| **Fragmentation of the actin distribution:**<br>**Fisher Information** |
| **Dynamic instability of the actin distribution:**<br>**Jensen Shannon Divergence** |

To verify that the mechanism underlying axon extension in the Abl KD and Abl OE conditions is fundamentally the same as that in wild type, we first queried the parameter set by performing a Principal Components Analysis (PCA) using the wild type dataset of growth cone measurements (Supplemental Figure 7). The first two principal components account for 47% of the total variance in the data set. PC1, which accounts for 29% of the variance, is dominated by contributions from morphological parameters including the number and length of protrusions along the axon, while PC2 (18% of the variance) is dominated by parameters that measure how the actin distribution is organized. We applied the two eigenvectors to the Abl KD and Abl OE data sets to compare how similar the global relationships among parameters in the altered-Abl conditions are to those in wild type (Figure 4A). A substantial number of data points from the Abl perturbed conditions occupied the same PC1 vs. PC2 parameter space as did the wild type data set. Crucially, however, the data points from Abl KD and Abl OE that did not overlap wild type did not just spread out isotropically, nor did they segregate into a discrete, separate domain of parameter space. Instead, they formed a restricted distribution along a single vector emanating from the cloud of wild type data. Stated otherwise, the ratio of the values of any pair of principal components in the Abl-perturbed conditions varied along a simple, linear relationship relative to their values in wild type, even for the data points with the most severely altered absolute values. This pattern was observed in all pairwise comparisons among the first 4 principal components, which account for 71% of the variance in the data set, thus revealing consistent quantitative relationships among the growth cone parameters that produce morphology and motility in wildtype, Abl KD and Abl OE. This verifies that the fundamental relationships among morphological and cytoskeletal properties of wild type growth cones are maintained in the Abl KD and OE conditions, even for the most severely affected data points in our dataset, and therefore validates the use of the altered Abl conditions to interrogate wild type growth cone dynamics.

**Figure 7:**
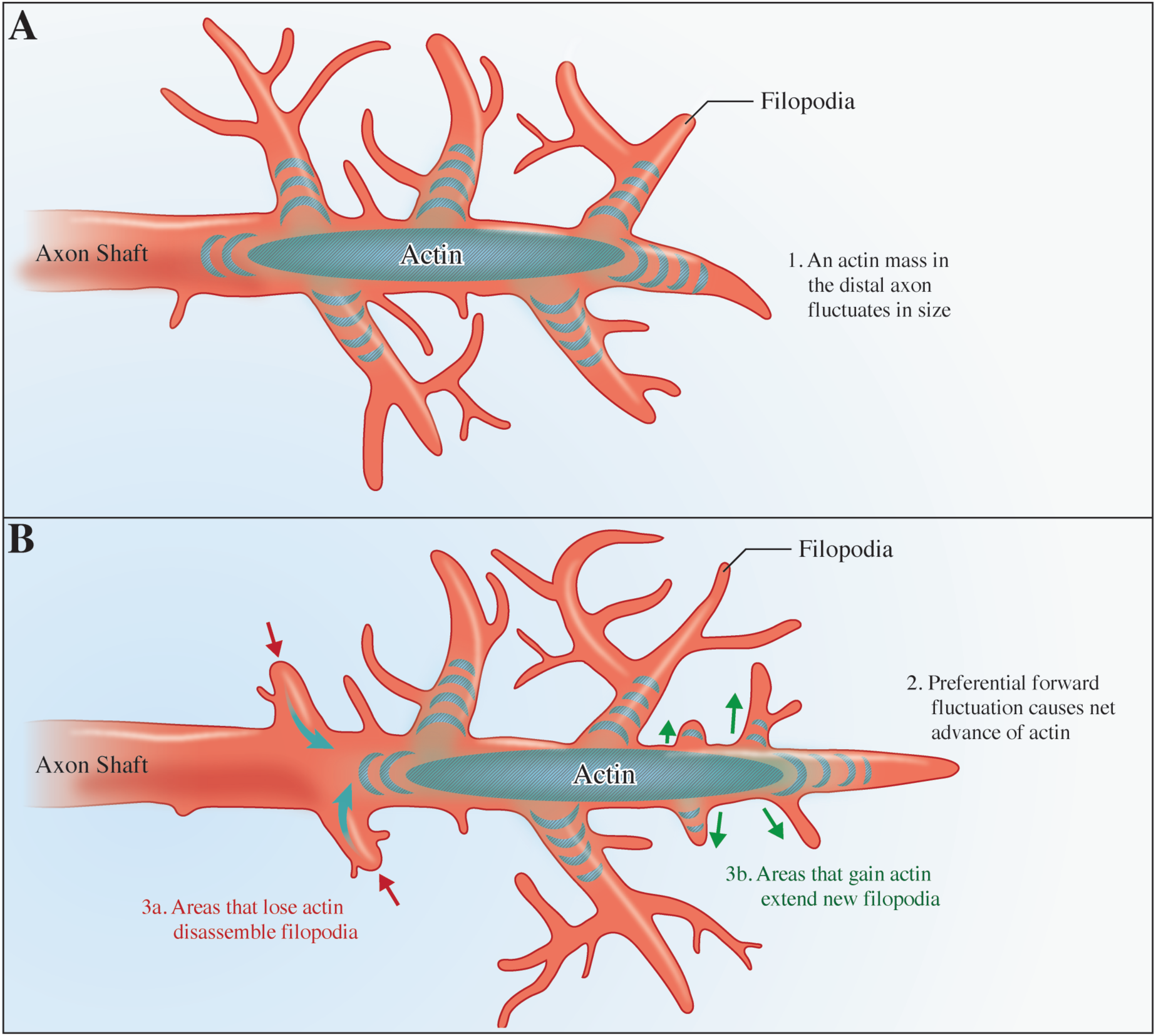
Model of actin-driven axon extension and consolidation. A) Accumulated actin in the distal axon fluctuates in size, transiently invading growth cone protrusions. B) Selective accumulation of actin in a more distal axonal protrusion advances the distribution, and triggers the emergence of nascent protrusions in that region, as well as the disassembly of protrusions in proximal regions no longer bearing enhanced levels of accumulated actin.

### Abl perturbation has large effects on the actin distribution and relatively smaller effects on growth cone morphology

Comparison of parameters across all three genotypes revealed that the actin distribution was far more sensitive to changes in Abl levels than were the morphological parameters of the axon, consistent with the hypothesis that actin remodeling may be the more direct mechanistic target of Abl-dependent signaling, upstream of changes in axonal morphology. Abl perturbation affects many growth cone parameters, but to varying degrees. For example, the mean length of the actin peak is decreased in the knockdown condition, and increased by Abl OE, yet is unaffected by expression of Dicer 2 alone (mean ±SEM, 15.16 *µ*m ± 0.19 in control vs. 14.0 *µ*m ± 0.27 in KD; p = 0.0051, 15.83 *µ*m ± 0.24 in Dicer 2; N.S., and 16.72 *µ*m ± 0.32 in OE; p = 0.0002; One – way ANOVA) (Figure 4B). Similarly, the length of the protrusion peak in Abl KD is smaller than wildtype (15.9*µ*m ± 0.21 in control vs. 14.5*µ*m ± 0.18 in KD; p<0.05) and is larger in Abl OE (18.9*µ*m ± 0.27; p< 0.001) (Figure 4C). To determine which aspects of TSM1 architecture were most affected by Abl, we calculated the Z-score for each parameter relative to the wildtype mean. While disruption of Abl signaling does not shift the mean value of any morphological feature by more than 0.7 standard deviations from its wildtype mean, actin parameter values are shifted by as much as 2 – 11 standard deviations away from their respective control means (Figure 4D). As Abl perturbation causes substantially larger effects on the distribution of actin in the axon as compared to axonal morphology, we infer that the actin distribution is the more sensitive target of Abl signaling.

### Net forward motion of the actin distribution arises from biased stochastic fluctuations of the distribution

In light of the observation that the shape of the actin distribution is a sensitive target of guidance signaling downstream of Abl, and that the pattern of actin redistribution predicts where future axonal protrusions will grow, we next examined the evolution of the actin distribution itself over time in each trajectory. We found that anterograde translocation of the actin peak was not consistent. Rather, both the maximum of the actin peak, and its midpoint, displayed extensive, seemingly stochastic, fluctuations proximo-distally along the axon, but with a small anterograde bias that resulted, over time, in net forward motion of the actin peak (Figure 5B). This bias in the stochastic motion of these positions is evident upon tracking the positions of the front, rear and maximum of the actin distribution over time for any single trajectory (Figure 5A). Further inspection of actin redistribution patterns revealed that the position of the actin maximum largely fluctuates at each time step within the domain defined as the actin peak (i.e. the window defined by the leading and trailing square root of the second moment of the actin peak; 77% of time steps in wild type (251/327). Consistent with this, we found that a broader actin peak allowed larger excursions of the maximum and midpoint, while narrow actin peaks are correlated with relatively short fluctuations of the position of the actin max (Pearson r = .24, p < 0.0001) (Figure 5C). The net effect is that the actin distribution displays an ‘inchworm’ style of motion, with the front of the actin peak advancing over time, and the rear catching-up (Figure 2G).

### Actin accumulation is fragmented and advance of the distribution is unpredictable in Abl-perturbed backgrounds

The most striking consequence of Abl deregulation in individual images is fragmentation of the actin distribution (Figure 6A - D). We quantified this phenotype using a measure from information theory called the Fisher Information Content (FI), which measures the coherence of the actin distribution, including both short-range fluctuations of intensity (<1*µ*m spatial scale) and mid – long range fragmentation of actin (2-20 *µ*m scale). We find that the actin distribution profiles from Abl perturbed axons are vastly fragmented as compared to controls (3.87 ± 0.14 in control vs. 32.67 ± 1.76 in Abl KD and 18.54 ± 1.40 in Abl OE; mean ±SEM, p<0.001 in each comparison) This suggests that one effect of Abl is to regulate the coherence of the actin distribution in the axon.

Next, we compared the actin distribution dynamics in the wildtype condition with the Abl perturbed conditions and found that the actin distribution evolves in an orderly way from one-time step to the next in wild type, but much less so when Abl is dysregulated (Figure 6E - 6H’, and Supplemental movie 2 - 4). The evolution of the actin distribution was quantified using another approach from information theory, called the Jensen Shannon Divergence (JSD), which calculates the dis-similarity (“divergence”) of the shape of the actin distribution between pairs of time points in a given trajectory. Thus, two distributions that are very similar have very low divergence (JSD close to 0), while two distributions that are very different have high divergence (JSD close to 1). In the control condition, successive time points of any single trajectory tended to have highly overlapped distributions, and thus low JSD values, and the overlap tended to decay in an orderly way as comparisons were made to more distant time points. In contrast, in the Abl KD and Abl OE conditions, the overlap between nearest neighbor time points was commonly as dissimilar as two distributions separated by multiple time steps in the wildtype condition, as indicated by increased mean JSD values (0.17 ± 0.0039 in the control vs. 0.38 ± 0.011 in the KD and 0.29 ± 0.011 in the OE; mean ±SEM p<0.001). These data demonstrate dynamic order in the temporal evolution of the actin distribution in each WT TSM1 trajectory, while in the Abl perturbed conditions, that same evolution is dynamically disordered and unpredictable.

### Actin disorganization correlates with morphological phenotypes in Abl perturbed backgrounds

While the strongest effects of Abl perturbation on single parameters were on those measuring aspects of the actin distribution, analysis of parameter interactions suggested that the critical function of Abl lies in the linkage of those actin parameters to their morphological consequences. This was revealed most clearly by the morphological correlates of the extremes of those actin values. Thus, for example, high fragmentation of the actin distribution was preferentially associated with the smallest growth cones in Abl KD (r = −0.14; p = 0.005), but it was associated with the largest GCs in Abl OE (r = 0.25; p < 0.0001) (Figure 6I). In the case of Abl KD, this reflected a population of very short growth cones with hypercondensed actin, that is, growth cones containing small, intensely-labelled foci of actin. Data presented above showed that actin step size, ie the distance the actin maximum (or midpoint) advances in any single time step, is correlated with the length of the actin peak (Figure 5C). In Abl KD, however, the growth cone often contracts beyond the lower limit observed in wild type, and the step size falls to nearly zero (Figure 6J). This is a genotype in which we observe growth cone stalling, and failure to form an axon. In contrast, in Abl OE, a genotype associated primarily with axon misrouting, maximal actin fragmentation is associated preferentially with the largest actin distributions, and in particular, with distributions broader than the upper limit typically observed in wild type (Figure 6K). These abnormally broad distributions often greatly exceed the distance over which the motions of individual actin molecules display correlation within a disordered actin network (decay length ∼11-16*µ*m; Ref 23). The potential molecular basis of these length-dependent changes in actin organization, and the consequences they are predicted to have for orderly axon growth, will be considered in detail in the Discussion.

## Discussion

What is a growth cone, and how does it extend an axon? We have shown here that a local accumulation of actin in the distal axon generates the zone of enhanced protrusive dynamics that defines the TSM1 growth cone. Over time, this actin bolus advances down the nascent axon, supporting formation of new filopodial protrusions from leading intervals now bearing enhanced actin levels, thus enabling extension of the axon. Protrusions left behind in the wake of actin advance are fated for disassembly, thus consolidating the proximal axon. Inchworm-like dynamics advance the actin mass with an anterograde bias that moves the distribution forward over time. Furthermore, our data demonstrate that the Abelson tyrosine kinase, a conserved regulator of cytoskeletal dynamics that signals downstream of many guidance cue receptors, coordinates the actin fluctuations that sum to produce the net forward motion of the distribution. Taken together, our data suggest that the fundamental function of Abl during axon guidance and extension is to modulate, in a probabilistic way, the fluctuations and the coherence of an advancing actin wave that directs the construction and consolidation of the growing axon in response to guidance cues (Model Figure 7).

Growth cone advance is often discussed by invoking deterministic, clutch-like adhesive mechanisms that harness the mechanochemical properties of leading lamellipodia ^1,2,24^. For some neurons, particularly those extending axons on relatively rigid, highly adherent substrata, these models provide a plausible explanation for axon growth. However, TSM1 and many other axons look non-lamellar, particularly in compliant, 3D environments like those often encountered by pioneer axons *in vivo* ^6,7,25,26^. In these contexts, growth cones are often dominated by filopodial protrusions and seem to lack the large veil-like structures that we associate with the adhesive style of growth. Moreover, the growth of these axons appears to be accomplished by protrusion and selective stabilization of filopodia, rather than by traction forces applied to the lamellipodia-like veils between filopodia (O’Connor et al., 1990; Sabry et al., 1991). These contrasting styles of axon growth are reminiscent of the dichotomy between cellular motility on rigid 2D substrata vs protrusive cell motility in compliant 3D environments. They also add support to the growing notion that the mechanisms of motility employed by neurons are not fixed, but are instead dependent on the context of their environment.

We now find that the filopodial-dominated TSM1 growth cone extends its axon within the intact *Drosophila* wing by regulated advance of an actin distribution, using a protrusive mechanism of motility that is probabilistic rather than deterministic, and is based on the statistical properties of disordered actin networks. Forward motion of the actin peak arises from a small spatial bias that is applied to the fluctuating actin distribution; forward motion of the axon terminus arises from preferential extension of filopodia from axon regions that now have high actin density and retraction of filopodia in regions of low actin density. In essence, the advancing actin peak directs processive axon growth by locally promoting assembly of potential axonal tracks, while the axon cannibalizes filopodia that lag behind the actin peak, and correspond to axonal paths that were not taken.

The key to such a mechanism for axon extension is that forward expansion of the actin distribution must be large enough to advance the actin bolus and produce net growth, yet constrained enough that the actin bolus remains coherent and the growth cone behaves as a single, unitary entity. We see here that balancing these two competing, yet related, requirements is the fundamental role of Abl tyrosine kinase signaling. It is well established that signaling by the Abl network must be maintained at an intermediate level of activity to support proper growth and guidance of axons (Grevengoed et al., 2001; Grevengoed et al., 2003; Liebel et al., 2003; Koleske et al., 1998; Moresco et al., 2005). Our data suggest that this intermediate level is crucial for two reasons. First, it minimizes the disorder of the accumulated actin distribution, as assayed both instantaneously, by the Fisher Information of the actin profile at each time step, and also dynamically, by the Jensen-Shannon analysis of the evolution of the distribution between time steps. Second, we also found that Abl regulates the width of the actin peak. This is seen most clearly in the Abl perturbations, where reducing Abl promotes condensation of the actin mass while increasing Abl promotes expansion of the peak. Specifically, Abl knockdown causes the actin bolus to hyper-condense and fragment into small, tightly-packed foci that show reduced spatial motion (Figure 4 and 6). Since expansion of the distribution provides the forward motion required for ‘inchworming’, it is plausible that failure of that expansion in the Abl KD condition is responsible for the growth cone stalling observed in this genotype. Conversely, overexpression of Abl causes the actin distribution to expand, often to the degree that the peak exceeds the characteristic decay length over which the motions of individual actin molecules remain correlated within disordered actin networks (∼11-16*µ*m) ^23^. This likely contributes to the preferential fragmentation we observe of the broadest actin peaks in the Abl OE condition, and plausibly could predispose to axonal misrouting and inappropriate axon branching in part by allowing different portions of the same growth cone to act independently. The two key functions of Abl, minimizing disorder and controlling distribution width, are closely intertwined; in the Abl KD condition the tendency is for the smallest growth cones to drive measures of disorder, while in Abl OE, the degree of actin disorder increases with expanding length of the actin peak (Fig 6I).

Our model of growth cone behavior is derived partly from the consequences of vigorous experimental perturbations of Abl, but our goal is to understand the consequences of modest modulation of Abl by guidance receptors. The principal component analysis, however, (Figure 4) demonstrates that the relationships among actin and morphological parameters in the Abl-manipulated experimental contexts are a faithful mirror of the relationships among those parameters in the wild type condition as well. Therefore, we infer that the more modest decreases and increases of Abl activity produced by guidance receptors in the course of wildtype TSM1 development modulate actin condensation and expansion similarly to what we see experimentally, albeit to a more measured extent. Hence, we would predict, for example, that a guidance cue that enhances Abl activity, such as Netrin acting through DCC/Frazzled ^27,28^ would promote expansion of the actin domain, and thus bias growth, in the direction of greater cue concentration, while local accumulation of a cue that leads to reduced Abl activity, such as Delta acting through the receptor Notch ^10,29,30^ would locally promote condensation of the actin, and therefore focus future proliferation of filopodia from the vicinity of that accumulation. Similarly, by this model it is straightforward to see why the subcellular localization that has been observed for some guidance receptors within specific subregions of the growth cone (V. Castellani, pers comm, Sept 2018) might be essential for them to promote either axon extension or retraction, respectively, or how a single receptor could be switched between attraction and repulsion simply by changing the polarity of its distribution in the growth cone.

A crucial mechanistic question raised by our experiments is how varying the level of Abl activity produces changes in the width of the actin peak. We have shown recently that the structure of the Abl signaling network intrinsically causes it to modulate the ratio of linear vs branched actin within the cell ^17^. Thus, activating Abl suppresses Enabled, a factor that extends linear actin polymers, but stimulates the Trio, Rac1, WAVE/SCAR, Arp2,3 axis that leads to actin branching. Suppressing Abl activity has the opposite pair of effects. Remarkably, a number of recent experimental and computational analyses have demonstrated that simply altering the ratio of linear vs branched actin in a disordered network is sufficient to alter network dimensions in just the way we observe in TSM1^23,31,32^. Thus, manipulations that produce increased network crosslinking, such as a higher proportion of branched actin, tend to expand a moderately cross-linked actin network, while increasing the proportion of unbranched, linear actin favors coalescence of the distribution. These consequences of actin network modulation are consistent with data presented here which show that the Abl KD condition causes growth cones with short, hypercondensed actin distributions to be overrepresented, while in the Abl OE condition, growth cones with longer, extended actin distributions are overrepresented (Figure 4 and supplemental Figure 8). Therefore, it is possible that the effect of Abl on the actin distribution, and therefore on morphological growth and guidance, could be accounted for simply by the demonstrated biochemical effects of Abl on pathways leading to branched vs linear actin. This conjecture will be investigated experimentally in future studies.

It is well-established that Abl has other roles aside from regulating actin extension and branching, in particular regulation of the microtubule plus-end tracking protein Orbit ^33^. Our data do not exclude the idea, for example, that Abl-dependent interaction of actin with microtubules could contribute to the forward-bias of actin fluctuations, perhaps by providing directionality. Additionally, the Abl network may also influence axon extension and guidance through more direct effects on morphological features of the growth cone ^27,28^. Finally, in this analysis, we have tracked the bulk distribution of actin. The motions of individual actin molecules remain unknown. Evolution of the actin distribution presumably incorporates actin polymerization, depolymerization, branching, crosslinking, diffusion, myosin contractility, active transport via microtubules, and bulk flow of actin from the cell body, among other contributors. Future experiments will aim to separate the individual contributions of these processes.

The mechanism of axon growth we observe here is energetically expensive, with only ∼5-15% of the total back-and-forth motion of the actin peak being captured in net advance of the actin mass. Considered in context, however, this mechanism allows an extremely efficient use of guidance information. The fluctuations of the actin peak cause it to repetitively sample leading and trailing positions along the axon multiple times before moving irrevocably along its trajectory. We propose that this is useful, and probably essential, for responding accurately to the shallow, often noisy gradients of individual guidance cues, which are typically presented in complex combinations in the substratum. In this sense, the mechanism we have found is most akin to the mechanism of bacterial chemotaxis, in which external attractants and repellants provide only a subtle spatial bias to essentially stochastic fluctuations of the motility machinery, relying on a ‘random walk with a ratchet’ to produce net guidance ^34,35^. Additionally, we note that these actin fluctuations survey all growth cone protrusions – off-axis lateral projections as well as on-axis extensions of the axon shaft– thereby allowing growth cone turning to derive from precisely the same machinery as does linear extension of the axon.

The wave-like, anterograde propagation of an actin distribution, regulated by the conserved Abelson tyrosine kinase, which probabilistically localizes the site of filopodial dynamics, represents a novel mechanism of motility that drives the extension and guidance of the TSM1 axon in the intact *Drosophila* wing. This style of growth expands our understanding of the fundamental mechanisms used by neurons to build neural circuits and reveals a novel molecular and cellular mechanism for Abl, one of the central regulators of cell morphology and neural wiring.

## Materials and Methods

### *Drosophila* stocks

*Neuralized-Gal4, Abl* knock down (KD) (*UAS-Abl-RNAi*), *Abl* overexpression (OE) (*UAS-Abl*), *UAS Dicer2*, and the *abl*^*4*^ allele have been described previously ^17,18^. *UAS-lifeactGFP*, and *UAS-CD4+ tandem tomato* stocks were obtained from the Bloomington *Drosophila* stock center.

### TSM1 axon growth and guidance phenotypes

White pre-pupae marked at the start of pupariation in each genotype were aged 7.5 – 8hrs at 25°C. Aged pupae were dissected in fresh culture media (CM) composed of Schneider’s *Drosophila* media (Life Technologies) supplemented with 10% fetal bovine serum (Gibco). Wings were removed and placed in a 35mm MatTek glass bottom microwell petri dish ventral side down and a halved circular cover slip was placed on their dorsal surface. Discs were slightly compressed to immobilize and flatten the tissue. Inert, compressible clay was placed between the bottom interior surface of the MatTek dish and the top cover glass to prevent over compression of the wings and to adhere the top coverslip to the bottom of the culture chamber (Supplemental Figure 1). 5 ml of CM was added to the dish after the tissue was secured in position. To assay terminal axonal phenotypes, wing discs were aged for ∼16 hours at 25°C and microscopy was performed on a Zeiss NLO 880 w/Airy scan confocal microscope using a 25X 1.4 NA water immersion objective. Non-wildtype TSM1 axons assumed one of four distinguishable phenotypes: no axon extended, axonal stall, guidance defect, or aberrant collateral branches. For simplicity, all non-wildtype phenotypes were scored as aberrant.

### Pharmacological inhibition of Abl

A 100mg tablet of Imatinib (Novartis, obtained from the NIH Clinical Center Pharmacy) was dissolved in 20 ml of Schneider’s *Drosophila* medium producing a 10mM stock, and aliquots were stored at −20C. 5ul of 10mM Imatinib/Schneider’s suspension was added to 4.995ml of CM for a final experimental concentration of 10μM. 5ml of Fresh 10μM Imatinib CM was added to wildtype wing discs mounted as described above. Cultures were aged and imaged as described above.

### Growth cone time-lapse imaging

Wing discs for the control, Abl-KD and Abl-OE conditions were prepared and mounted as described above for scoring aberrant end-stage phenotypes. Live-imaging was performed immediately after mounting at 25°C using an inverted Zeiss Axio Observer Z1 confocal microscope fitted with a Yokogawa Spinning Disk (SD) module and a temperature-controlled stage. Z-stacks at 0.8um spacing were collected from individual TSM1 axons every 3 minutes using a 63X 1.2NA water immersion objective. (Note that laser power was set at a minimal value to mitigate photodamage and ensure image intensities were not saturated.)

### Segmentation of the axon and quantification of growth cone parameters

Neuronal z stacks were stereoscopically reconstructed in 4-dimensions (x, y, z, and time) using IMARIS (Bitplane, version 8.0.2) and segmented using the semi-automatic ‘filament tracer’ function. The axonal backbone and protrusions of TSM1 were identified and traced using the membrane localized CD4+ tandem tomato signal; false positives were removed manually, and untraced protrusions were added manually. Tracings were exported as inventor files (.iv) and converted to the standard SWC file format using a MiPav plugin. Morphological parameters extracted from these segmented images included the number of protrusions, length of each protrusion, position of individual protrusions along the axon, and protrusion branch order among others (for a full list of measured parameters, see Table 1)

Custom scripts in Mathematica software were written to identify computationally the position of highest filopodial density along the axon by using a sliding-window method to sum the length and number of protrusions within a 5μm window that advanced 1um/per step along the segmented axon. We varied window size from 1-10 μms and empirically found peak protrusion density assignment was insensitive to the size of the window (data not shown). We then calculated the square root of the second moment about the peak of protrusion density to determine the length of the protrusive zone, both separately for the portions of the distribution leading and trailing the peak position, and also globally to measure the length of the entire distribution.

### Actin distribution measurements

SWC files from a complete trajectory of TSM1 growth and the corresponding 4D Z stacks, (x, y, z, time) converted to Nikon image cytometry standard (ICS) format in Imaris, were loaded into two MiPav plugins, PluginDrosophilaCreatesSWC and PlugIn3DSWCStats to extract the actin distributions from each time point. Plugin code is available in MiPav, but its function is described below.

In brief, image intensity is calculated by summing the actin intensity within sequential frustums that encompass the axon. First, the MiPav plugin performs a background subtraction on each image. A 3D probability map of intensity values is then generated to determine the boundaries of the actin signal. A circle is then expanded from each coordinate in the SWC tracing to determine the radius of each frustum. The radius is normal to the SWC axis, and is bounded by the background of the image in the probability map. Summed intensity values from non-overlapping frustums are then reported as a function of position along the axonal tracing. Actin peak position was identified by sliding window and the square root of the second moment of the distribution was calculated just as for the equivalent measurements of filopodial density.

### Fisher Information

Fragmentation of the actin distribution was quantified by the Fisher Information (FI), which is a measure of the amount of information that can be specified by the shape of a distribution. The FI for each actin distribution profile was calculated according to the formula:

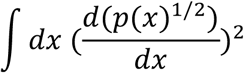

using the trapezoid approximation to the integral, where p(x) is the distribution of actin intensity (x) values for each actin profile. In this context, FI yields a quantity representative of the fragmentation of the actin distribution.

### Jensen Shannon Divergence

The Jensen-Shannon divergence (JSD) was used to quantify the divergence, or overlap, between actin distributions at different time steps of any single trajectory according to the formula:

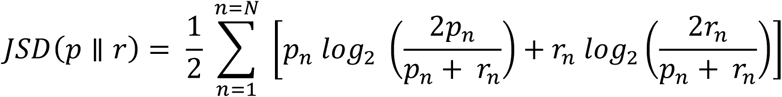

where p and r are two probability distributions, i.e. actin distributions, Since the JSD does not satisfy the triangle inequality, for the principal component analysis in this report we used the square root of the JSD, the Jensen –Shannon metric (JSM) which can be used as a metric.

### Quantification of protrusions in response to actin distribution translocation

To quantify the effect that actin advance has on local axonal protrusion, we summed the length of protrusions in a 10*µ*m interval surrounding the midpoint of the actin distribution at the start of a 20*µ*m translocation, and in a 10*µ*m interval surrounding the endpoint of the translocation (MATLAB).

### Statistics and reproducibility

All statistical tests, and their respective parameters, that were performed in this manuscript are reported in text and figure legends above.

### Data and Code availability

Numerical data for all figures is included in Supplemental datasheet 1. MiPav plugin code for extraction of actin intensity profiles has been incorporated and released in the publicly available NIH image analysis package package, MiPav. Matlab and Mathematica scripts will be made available upon publication by uploading to the publicly accessible NIH website: https://data.ninds.nih.gov

## Acknowledgements

We wish to thank all the members of our lab for their advice and assistance during the course of these experiments, particularly Kate O’Neill for Matlab expertise. We would also particularly like to thank Chi-Hon Lee, Sally Moody, Clare Waterman, Bob Fischer, Chun-Yuan Ting, Lenny Campanello, Wolfgang Losert and Garyk Papoian for their many helpful suggestions, and Laura Alto, Jon Terman and Ken Yamada for comments on the manuscript. Additionally, we thank Mike Murrell and Ian Linsmeier for sharing their data and ideas about the motions of actin molecules, and Valerie Castellani for sharing unpublished data on fine-scale localization of guidance receptors in the growth cone. Many *Drosophila* stocks were provided by the Bloomington *Drosophila* Stock Center. These experiments were supported in part by the Basic Neuroscience Program of the NINDS Intramural Research Program (Z01-NS003013 to EG). PGM, VW and EMcC were supported by the Intramural Research Program of NIH, CIT, and SW was supported by the NHGRI, NIH. RK was supported in part by a DBT Ramalingaswami re-entry fellowship from the Government of India.

---

**Supplementary Figure 1:**
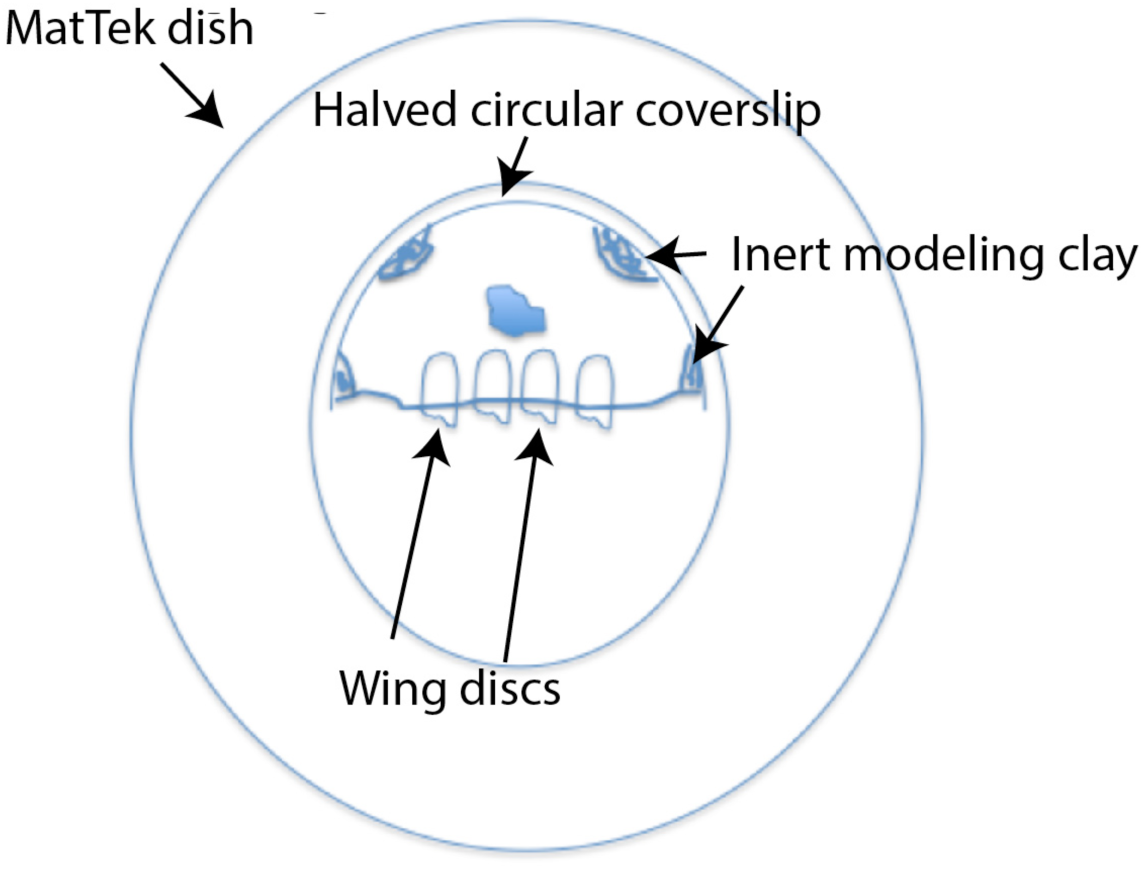
Mount schematic for culturing and imaging TSM1

**Supplementary Figure 2:**
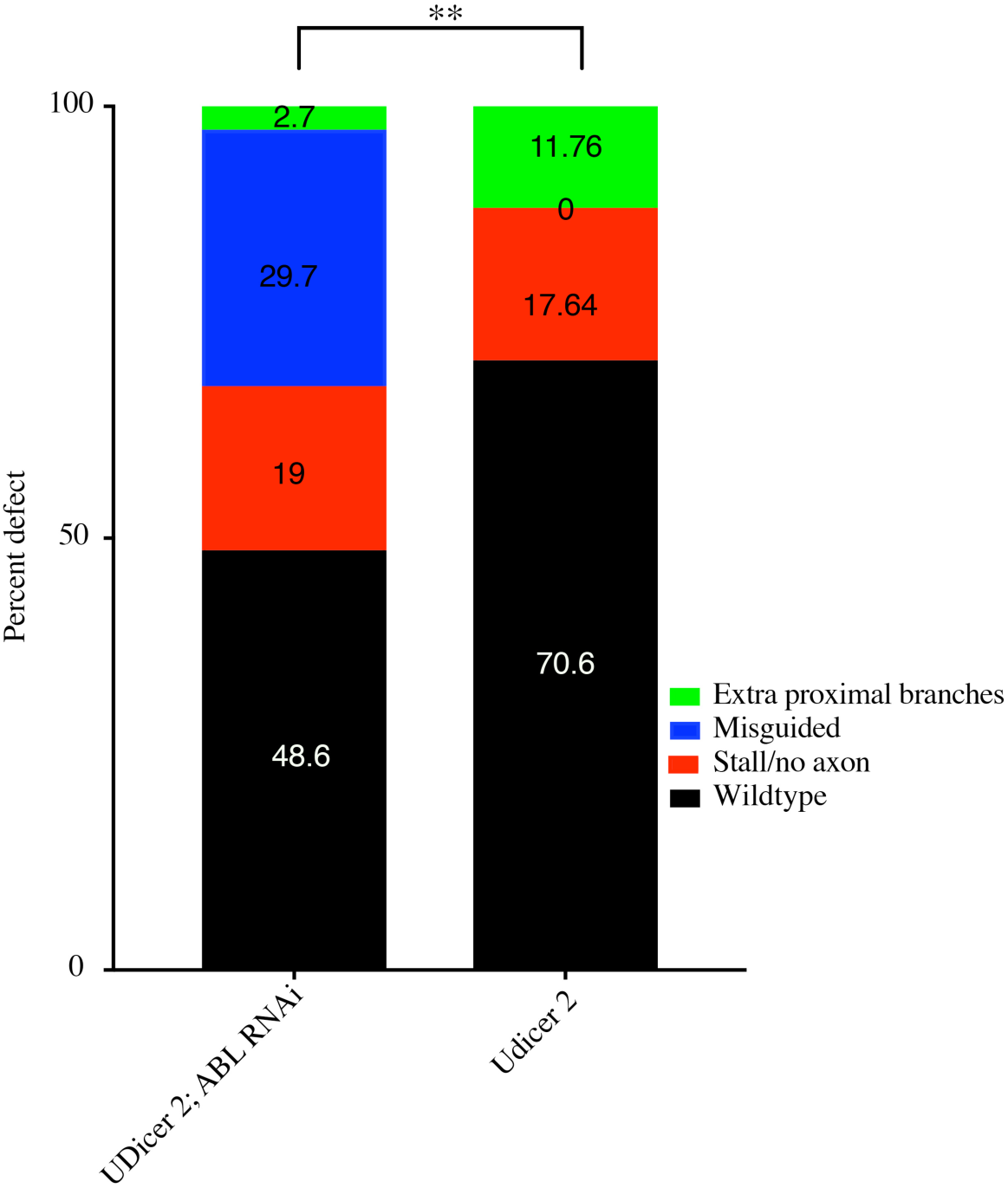
Abl KD enhanced by Dicer2 causes a different spectrum of defects than that caused by Dicer 2 alone. Chi-square; **p = 0.0035.

**Supplementary Figure 3:**
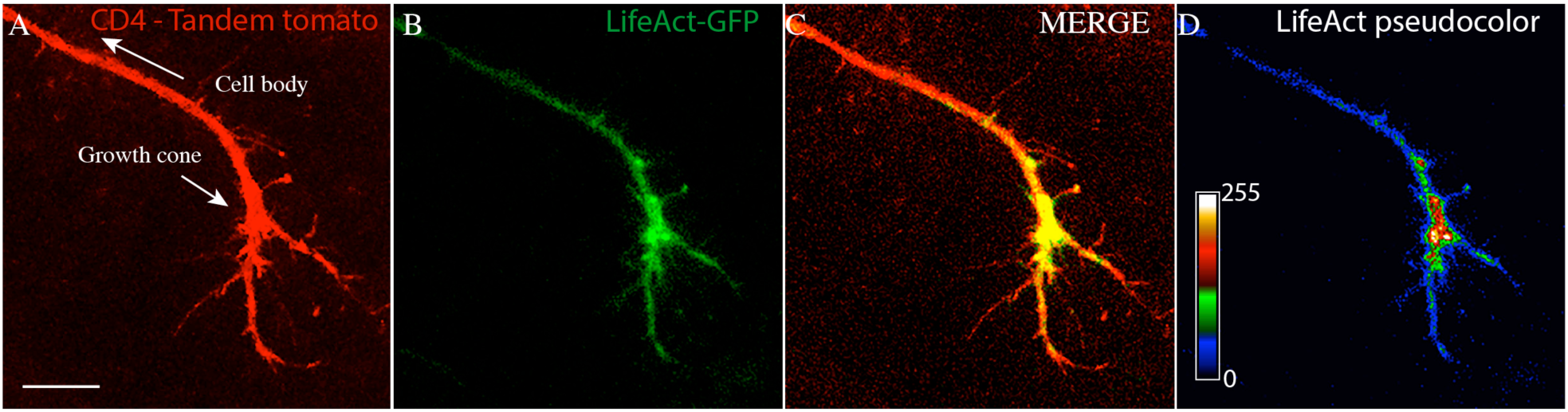
Actin accumulates at the site of the TSM1 growth cone. A) A max intensity projection of the membrane localized tandem tomato signal is presented alongside B), a summation projection of the lifeactGFP signal and C), a merge of both channels. The lifeactGFP summation projection is pseudocolored in D) to detail local GFP intensity values. Fiji/ImageJ was used for all image processing in this gallery.

**Supplementary Figure 4:**
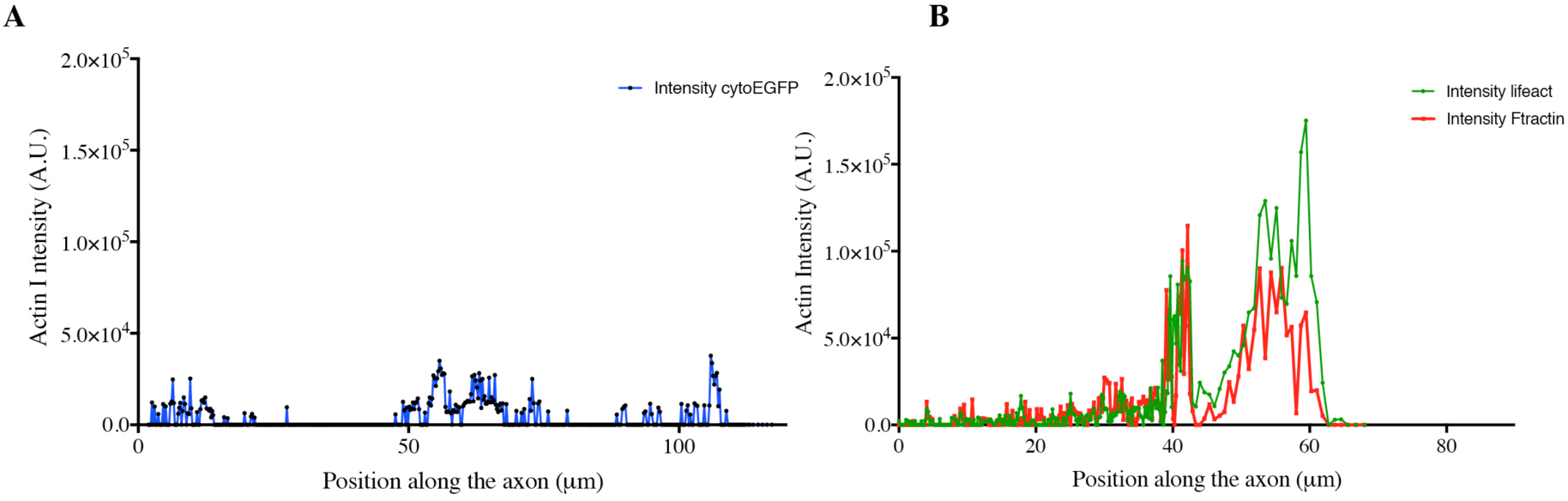
Different actin markers, but not cytoplasmic eGFP, accumulate in a distal mass in the TSM1 axon. A representative actin distribution extracted from a TSM1 axon expressing cytoplasmic eGFP. An overlay of two actin distributions from a single TSM1 axon expressing both the lifeactGFP and F-tractin transgenes. Each plot is representative of n = 5 independent trajectories, with > 15 time points per trajectory.

**Supplementary Figure 5:**
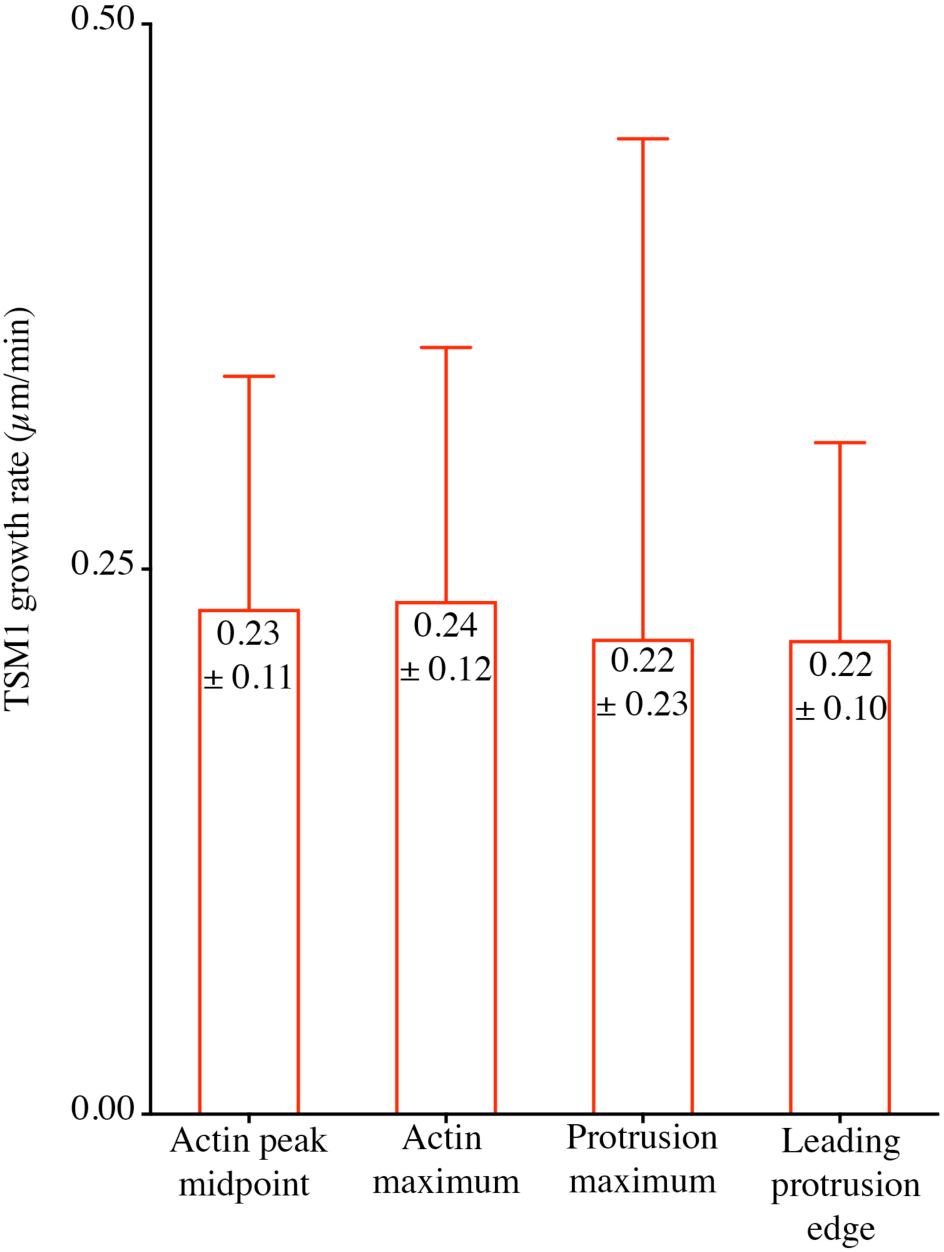
TSM1 growth rate. The indicated features of the TSM1 growth cone were tracked independently over time. Mean ± SEM are reported. Measures of growth rate are not significantly different as analyzed by ANOVA and paired T-test.

**Supplementary Figure 6:**
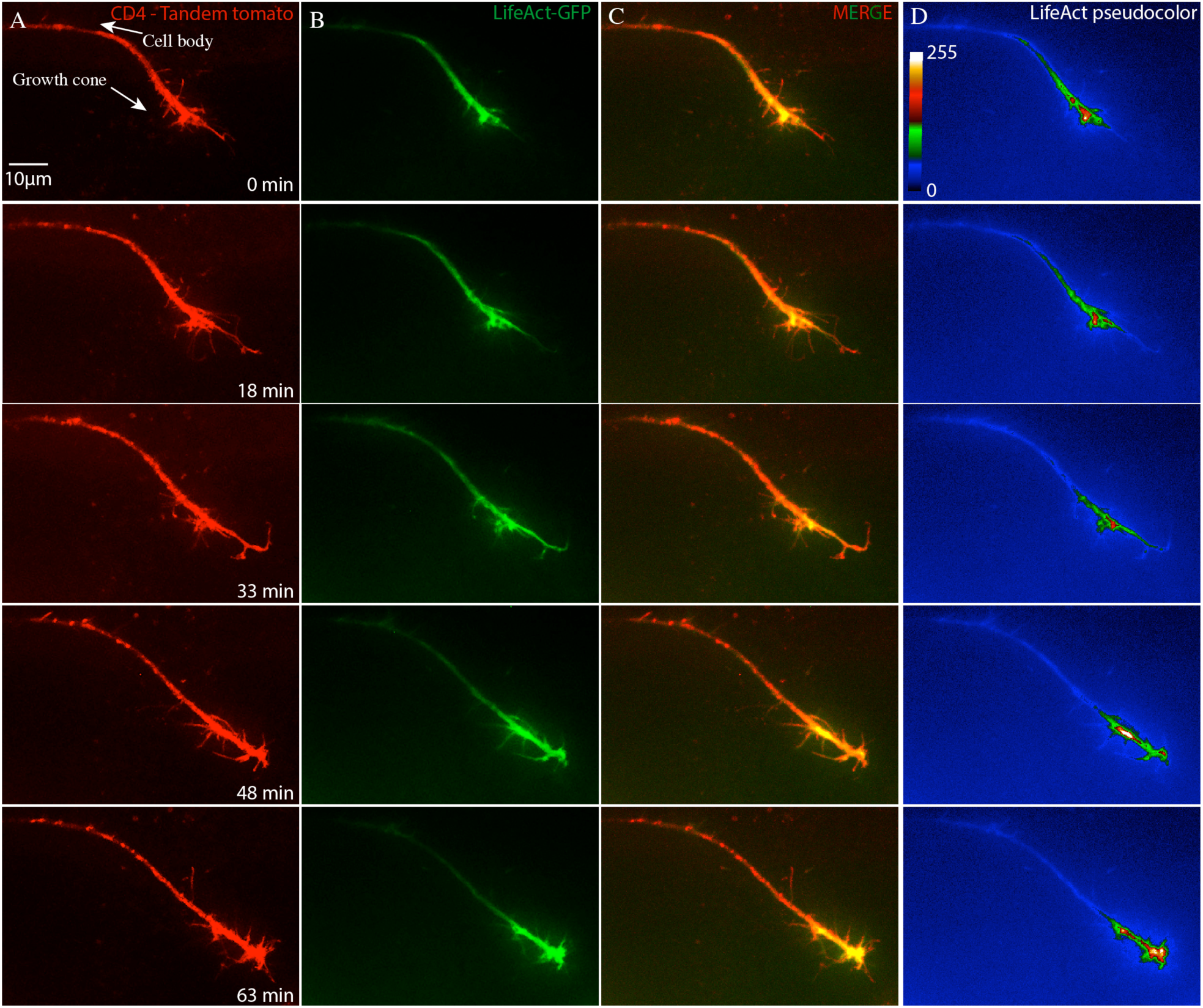
Actin accumulation advances the TSM1 growth cone. A time course gallery of (A) a max intensity projection of the membrane localized tandem tomato signal, (B) a summation projection of the lifeactGFP signal, (C) a merge of the membrane and actin channels, and (D), A pseudocolored lifeact summation projection detailing local GFP intensity values. Fiji/ImageJ was used for all image processing in this gallery.

**Supplementary Figure 7:**
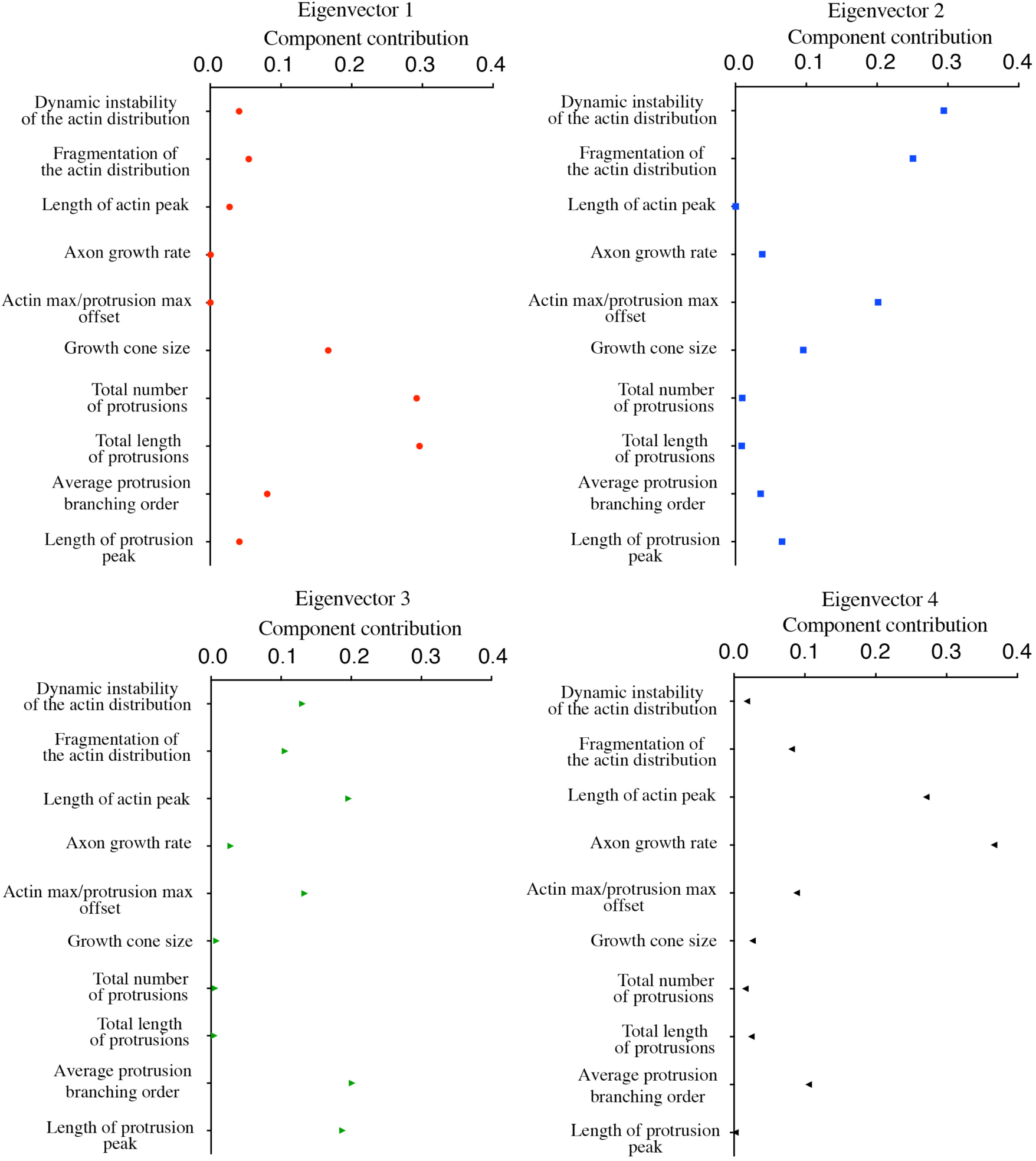
Parameter contributions to the first four principal components. Principal component analysis was performed using 10 measured variables that quantify aspects of TSM1 axonal growth. The first four eigenvectors account for ∼71% of the total variance in the data set.

**Supplementary Figure 8:**
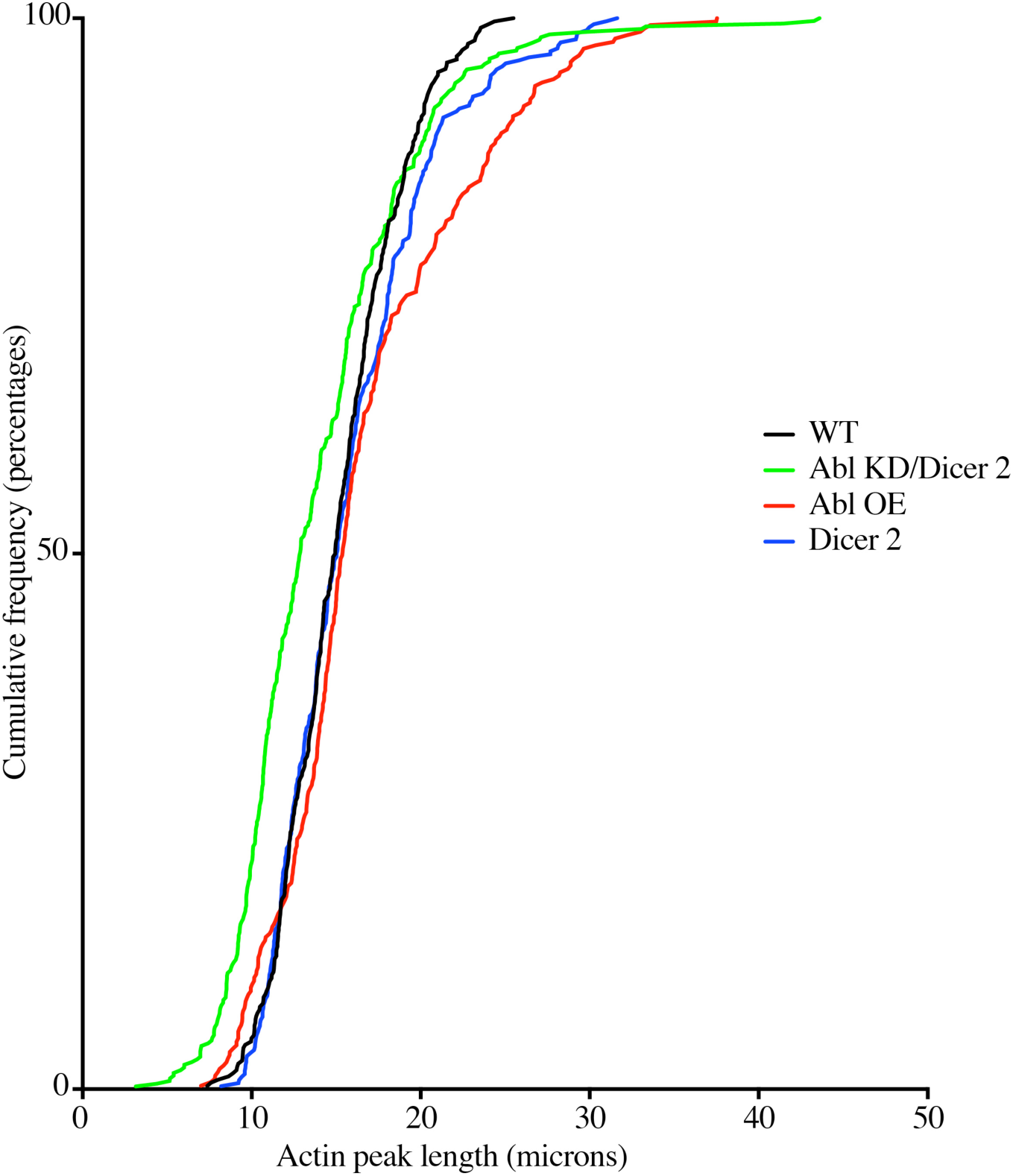
Cumulative frequency of actin distribution sizes. The lengths of the actin peak for each genotype are presented as cumulative frequency distributions to visualize the contribution of actin peaks of different lengths to each data set.

